# Terminating contamination: large-scale search identifies more than 2,000,000 contaminated entries in GenBank

**DOI:** 10.1101/2020.01.26.920173

**Authors:** Martin Steinegger, Steven L Salzberg

## Abstract

Metagenomic sequencing allows researchers to investigate organisms sampled from their native environments by sequencing their DNA directly, and then quantifying the abundance and taxonomic composition of the organisms thus captured. However, these types of analyses are sensitive to contamination in public databases caused by incorrectly labeled reference sequences. Here we describe Conterminator, an efficient method to detect and remove incorrectly labelled sequences by an exhaustive all-against-all sequence comparison. Our analysis reports contamination in 114,035 sequences and 2767 species in the NCBI Reference Sequence Database (RefSeq), 2,161,746 sequences and 6795 species in the GenBank database, and 14,132 protein sequences in the NR non-redundant protein database. Conterminator uncovers contamination in sequences spanning the whole range from draft genomes to “complete” model organism genomes. Our method, which scales linearly with input size, was able to process 3.3 terabytes of genomic sequence data in 12 days on a single 32-core compute node. We believe that Conterminator can become an important tool to ensure the quality of reference databases with particular importance for downstream metagenomic analyses. Source code (GPLv3): https://github.com/martin-steinegger/conterminator

## INTRODUCTION

The number of genomes in public and private repositories has been skyrocketing for at least the past decade, primarily due to the rapidly dropping costs of sequencing. The public genome database GenBank, which is regularly synchronized with the EMBL and DDBJ databases, has been doubling in size roughly every 18 months [1]. These genomics databases provide a vital worldwide resource that has been driving new findings in biotechnology and medicine for nearly three decades.

Draft genomes consisting of hundreds to thousands of unordered DNA sequence fragments represent a large fraction of the over 500,000 genomes stored in GenBank [2]. Some of these fragments contain foreign DNA due to contamination from reagents, laboratory materials, sample processing artifacts, or cross-contamination from multiplexed sequencing runs (Fig. 1a). These contaminating sequences may cause a variety of problems, including incorrect labels on sequences in metagenomic studies [3], faulty conclusions about horizontal gene transfer [4, 5], or poor annotation quality of genomes [6].

**FIG. 1.**
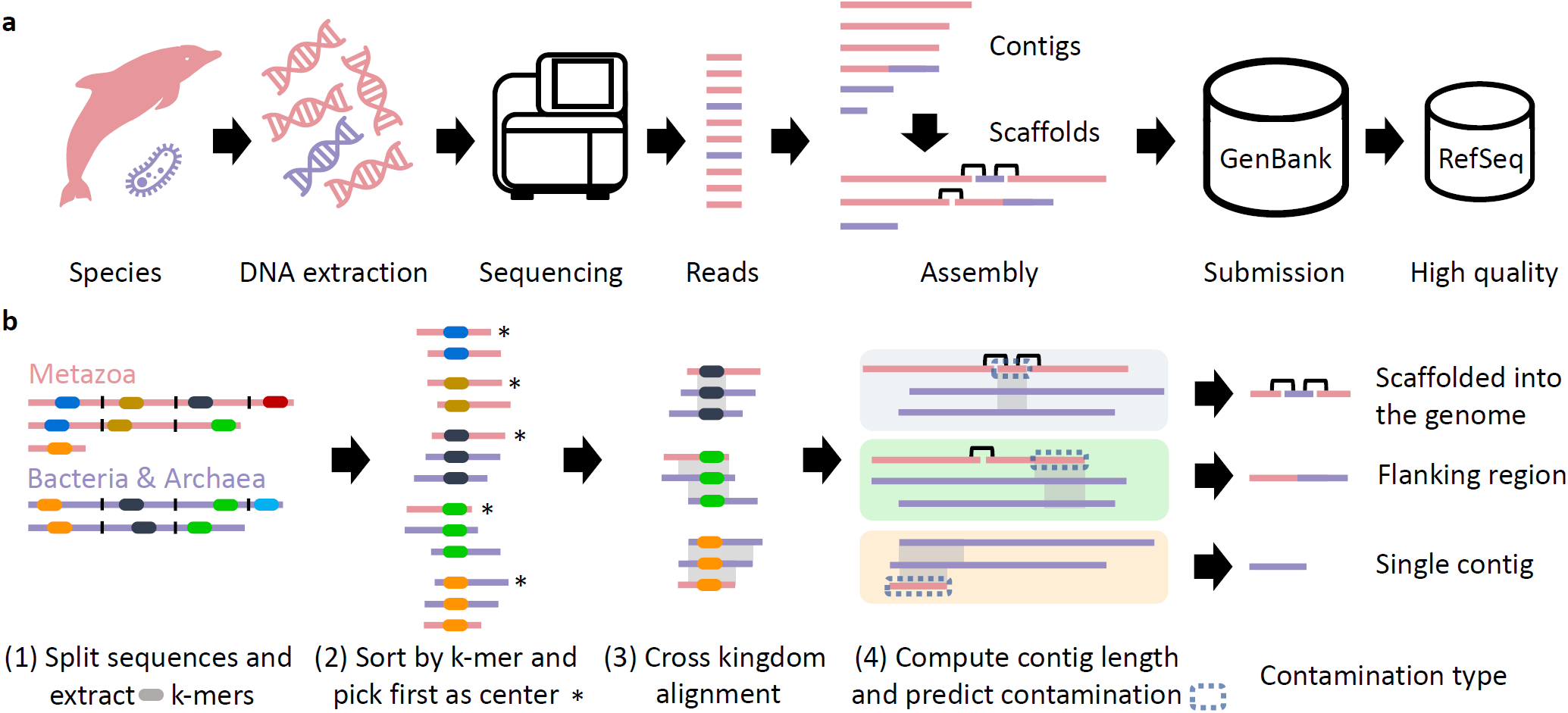
How contamination occurs and how Conterminator detects it. **a**, DNA extraction from an organism (red) is imperfect and often introduces contamination by other species (violet). DNA sequencing then generates short reads that are assembled into longer contigs. Contaminated DNA is typically assembled into separate, small contigs, but sometimes is erroneously included in the same contigs as DNA from source organism. Contigs may also be linked by scaffolding, which can produce scaffolds containing a mixture of different species. Final assemblies are submitted to GenBank, and higher-quality assemblies are entered in RefSeq. **b**, Conterminator detects contamination in proteins and nucleotide sequences across kingdoms; e.g., bacterial contaminants in plant genomes. The following describes the nucleotide contamination detection workflow. (1) We take taxonomically labeled input sequences and cut them into non-overlapping segments of length 1000 and extract a subset of *k*-mers. (2) We group the *k*-mers by sorting them and compute ungapped alignments between the first and all succeeding sequences per group. (3) We extract each region of the first sequence that has an alignment to other kingdoms that is longer than 100 amino acids (residues) with a sequence identity greater than 90 %. We perform an exhaustive alignment of the input sequence segments against the multi-kingdom regions. We offset the alignment’s start and end position to the respective coordinates in the input sequence. (4) We reconstruct contig lengths within scaffolds by searching for the scaffold breakpoints (indicated by N characters in the DNA sequence) on the left and right side from the alignment start and end position. We predict that contamination is present if an alignment hits a contig that is shorter than 20 kb that aligns to a different kingdom with an alignment length longer than 20 kb.

To combat the contamination issue, NCBI (the home of GenBank) applies two filtering protocols for detection of contaminated fragments. First, VecScreen [7] is used to detect synthetic sequences (vectors, adapters, linkers, primers, etc.), and second, BLAST alignments [8] against common contaminants identifies a broader array of contaminating sequences. Despite these filters, contamination still occurs, and its detection remains challenging [9, 10].

Because humans are always present in sequencing labs, *Homo sapiens* continues to be a major source of contamination for genome projects. Contaminating pieces of human DNA occasionally remain in published genomes [11] despite automated searches. A recent study, for example, showed that thousands of human DNA fragments can be found in draft bacterial genomes, and that many of these have been erroneously translated and annotated as proteins [10]. However, many other species also cause contamination [12–16]. Systematic approaches to detect contamination are limited by computational costs of comparing every submitted genome against all other known genomes. For example, a BLAST all-against-all comparison of the RefSeq database [17], which has a size of 1.5 Tb, would take ≈30,000 CPU years. Faster alignment method such as Minimap2 [18] or Bowtie2 [19] will take less time, but will still suffer from the quadratic complexity of this comparison.

We present Conterminator (Fig. 1b) a fast method for detecting contamination in nucleotide and protein databases by computing local alignments across taxonomic kingdoms. It utilizes the linear-time all-against-all comparison algorithm from Linclust [20] followed by exhaustive alignments using MMseqs2 [21]. This enables us to process huge nucleotide and protein sequence sets on a single server. We applied this method to quantify the current state of contamination in the nucleotide databases Genbank [1] and RefSeq [17], and in the comprehensive NR protein database [1].

## RESULTS

Figure 2 summarizes the contamination found by Conterminator in RefSeq (Fig. 2a,b) and GenBank (Fig. 2c,d). Processing the 1.5 and 3.3 terabytes in RefSeq and GenBank took 5 and 12 days on a single 32-core machine with 2 terabytes of main memory. Conterminator reported 114,035 and 2,161,746 contaminated sequences affecting 2767 species and 6795 in RefSeq and GenBank respectively. Identifiers of the contaminated sequences are available in Supplemental Files 1&2. In GenBank, over 95 % of contamination occurred in eukaryotic genomes. Eukaryotic genomes tend to be much more fragmented due to their larger genome sizes and higher repetitive content (as compared to prokaryotes), and many of the smaller contigs in eukaryotic genome assemblies suffer from contamination.

**FIG. 2.**
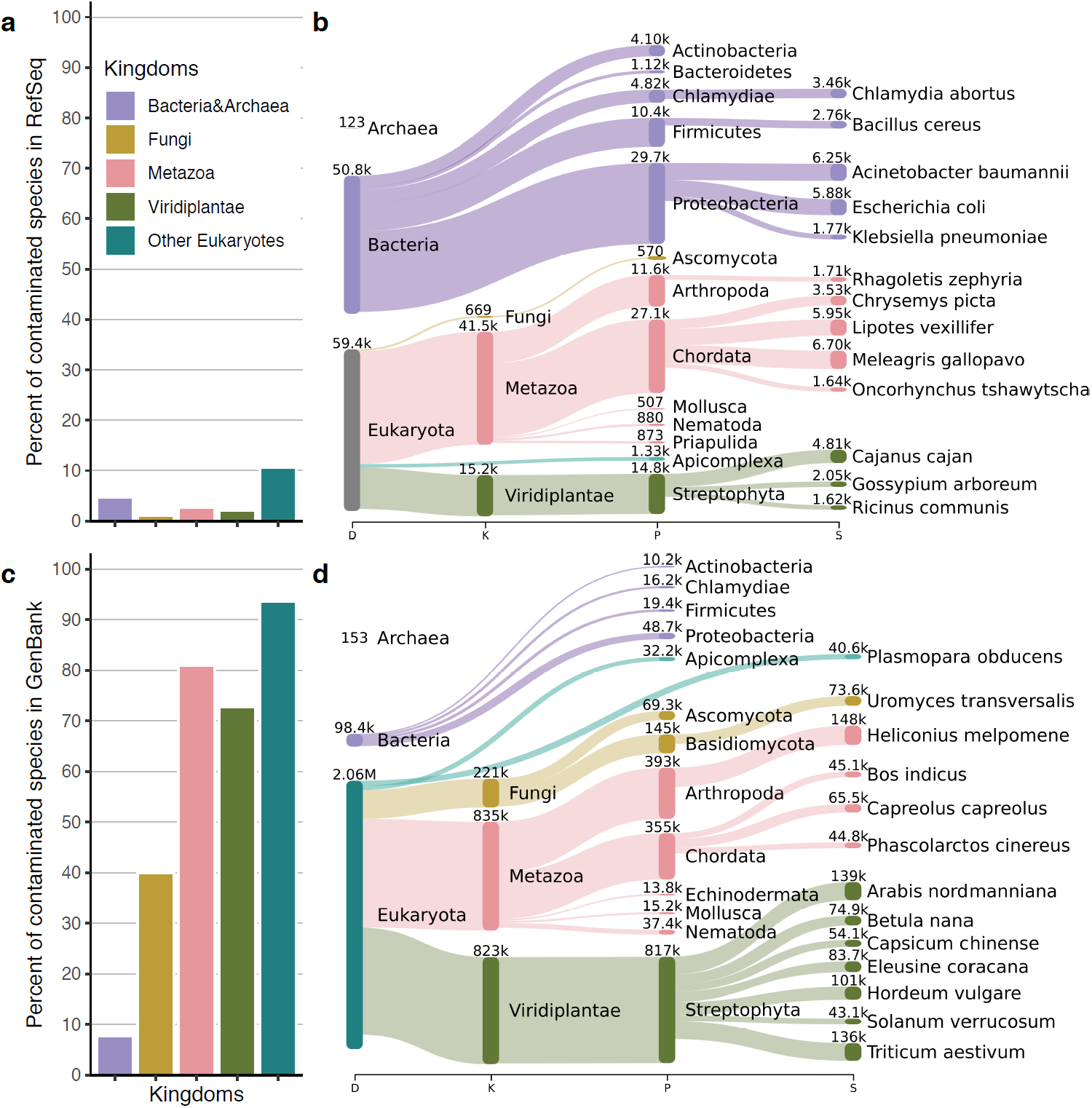
Results of contamination within the RefSeq. **a** Distribution of contaminated species in RefSeq across five kingdoms: Bacteria&Archaea (violet), Fungi (yellow), Metazoa (red), Viridiplantae (green) and other Eukaryotes (turquoise). **b** Sankey plot of the top 13 contaminated species in RefSeq. We show the taxonomic ranks domain, kingdom, phylum and species. Numbers show above each taxonomic node indicate the total number of contaminated sequences. The tree uses the same color code for kingdoms as in **a. c, d** Same as **a,b** but for GenBank.

In RefSeq only 52 % of the contamination occurred in eukaryotic genomes. One likely reason for this is the more stringent filters used to determine which Gen-Bank genomes are included in RefSeq; these filters reject genomes with very low contig sizes or genomes that were flagged as contaminated. The number of species identified as contaminants (i.e., the species causing contamination) in RefSeq was 2881 and in GenBank the number was 13,981. The leading contaminant species are *Homo sapiens, Saccharomyces cerevisiae, Stenotrophomonas maltophilia* and *Serratia marcescens* (see Supplementary Fig. 1).

### Contamination in high-quality genomes

We expected that well-studied model organisms would have the highest-quality genomes, and that these genomes would have very little, if any, contamination. We also expected very little contamination in finished microbial genomes. Therefore, we created a control set of high-quality genomes consisting of 928 genomes from FDA-ARGOS, a curated set of complete microbial genomes [22], plus genomes for model organisms *Saccharomyces cerevisiae, Danio rerio, Mus musculus, Drosophila melanogaster, Arabidopsis thaliana, Caenorhabditis elegans* and *Homo sapiens*. We searched for (presumably) false positive predictions by scanning our RefSeq results for contaminants in these high-quality genomes. Initially, our method did not report any contamination for any of these genomes, in part because by default it only reports contamination when the target sequence is shorter than 20 kb (see Methods). We then considered alignments to sequences longer than 20 kb in this high-quality genome set. In this additional scan, we found alignments between bacterial sequences and two eukaryotes: (1) *Acidithiobacillus thiooxidans* in *Homo sapiens* and (2) *E. coli* in *Caenorhabditis elegans*, shown in Figure 3.

**FIG. 3.**
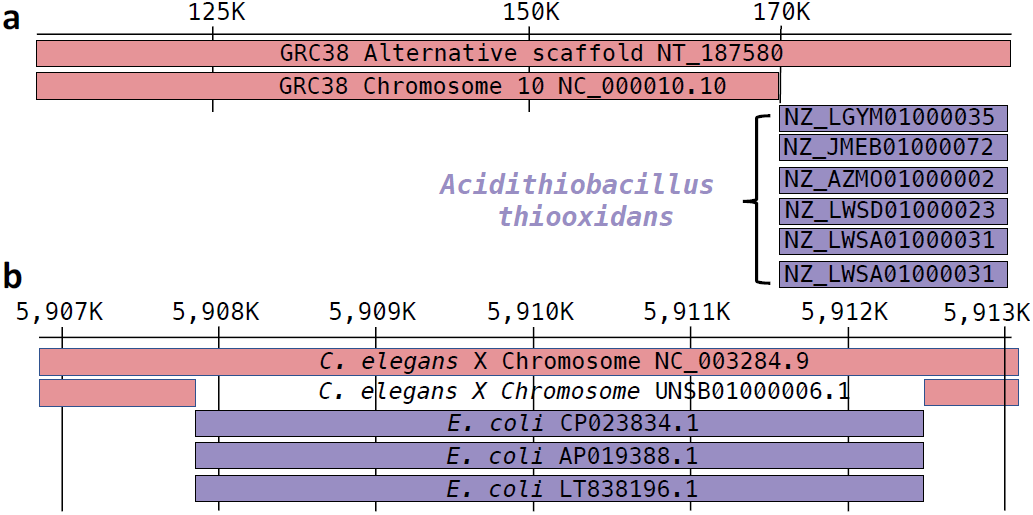
Contamination in the reference genomes of *Homo Sapiens* and *Caenorhabditis elegans*. **a** Alignment of *Homo sapiens* alternative scaffold `NT_187580` of chromosome 10 against RefSeq. Chromosome 10 (`NC_000011.10`) aligns with 100 % sequence identity from position 1 to 169,918. The remaining 18,397 residues of `NT_187580` align only to *Acidithiobacillus thiooxidans* at 98 % sequence identity. Shown are only 6 out of 15 alignments to *Acidithiobacillus thiooxidans*. **b** The X chromosome of *Caenorhabditis elegans* `NC_003284.9` aligns on the left and right flanking position around 5,907,856 until 5,912,458. *E. coli* genomes aligns from 5,907,856 to 5,912,087, a total of 4231 residues. Shown are only 3 out of 8199 alignments to *E. coli*.

### *A. thiooxidans* in human genomic sequence

The human reference genome (currently GRCh38) consists of chromosomal scaffolds, unplaced scaffolds, and “alternate” scaffolds. The last group is included in the reference genome to represent sequences that are divergent from the primary chromosome sequence. In NT 187580, an alternate scaffold on chromosome 10 in GRCh38.p13, we detected a sequence matching *Acidithiobacillus thiooxidans* that spans positions 169,917–188,315 of the human scaffold (Fig. 3a), which has a length of 188,315 base pairs. 15 different *A. thiooxidans* genomes align to the contaminated portion of the scaffold. The primary sequence of human chromosome 10 aligns perfectly from positions 1 to 169,918 on the alternate scaffold, but that alignment stops at the region that aligns to *A. thiooxidans*. Thus, the last ∼18 kb of this human alternative scaffold appears to be bacterial.

### *E. coli* in the *C. elegans* reference genome

Our method also detected a bacterial contaminant in the *C. elegans* reference genome, in chromosome X (GenBank accession `NC_003284.9`). A segment spanning positions 5,907,856–5,912,087 of the *C. elegans* sequence aligns perfectly to multiple strains of *E. coli* (Fig. 3b). To check whether this might be a false positive reported by our method, we downloaded the raw Illumina reads used for the assembly of this *C. elegans* strain (`SRR003808` and `SRR003809`) and aligned them against the chromosome X assembly (`NC_003284.9`) using Bowtie2 [19]. Only six reads (30 bp each) aligned in this region. In contrast, the average coverage over the rest of the chromosome was ∼99.8. This indicates that the *E. coli* sequence was indeed a contaminant. To corroborate the contamination further, we looked at a recent assembly of *C. elegans* that used a combination of long and short reads [23]. We aligned their assembly of chromosome X (GenBank accession UNSB01,000006.1) against the current reference, and found that in this newer assembly, the *E. coli* region is not present. This strongly suggests that the *C. elegans* reference genome contains a ∼4 kb insertion of *E. coli* contamination.

### *Meleagris gallopavo* genome cleanup

The most contaminated genome in RefSeq, on our initial scan, was the turkey genome, *Meleagris gallopavo* [24]. The contaminants included 6698 small, unplaced scaffolds with a total size of 2,655,271 bases. More than half of the contaminations were caused by *Achromobacter xylosoxidans* and *Serratia marcescens*. We contacted the original authors of that assembly to communicate our findings, and they subsequently removed all contaminated fragments, plus an additional 39,413 contigs that were shorter than 300 bp. The new version of the assembly, `Turkey_5.1`, has no contaminants and is available in GenBank as accession `GCA_000146605.4`.

### Proteins in contaminated RefSeq contigs

We detected that 19.4 % of the contaminated RefSeq contigs contain protein annotations and encode a total of 47,943 proteins. A previous study [10] reported 3437 spurious bacterial proteins that originate from human repeats that have contaminated bacterial genome assemblies. We aligned these sequences against our set using MMseqs2, enforcing a 80 % alignment coverage of the shorter sequence (`--comp-bias-corr 0 --mask 0 --cov-mode 5 -c 0.8`), and discovered that our set contains 62 % of the previously-reported proteins.

We clustered the proteins using MMseqs2 at a 95 % sequence identity, enforcing a bi-directional coverage of 95 % (`cluster --min-seq-id 0.95 -c 0.95`). This resulted in 3339 clusters that covered 12,494 sequences. The remaining sequences were singleton clusters. The largest cluster consists of 185 bacterial proteins, all of which are located on contigs shorter than 1 kb, and the proteins are widely spread among multiple phyla in the bacterial kingdom. Despite the long evolutionary distance, 166 of the sequences are 100 % identical to each other and the remaining are at least 95 % identical, suggesting that all of them represent contaminants (see Fig. 4). All contigs align to multiple positions in the domestic sheep *Ovis aries* genome; the fragments align to chromo-some 15 (`NC_040266.1`) with a sequence identity greater than 94 % with nearly complete coverage.

**FIG. 4.**
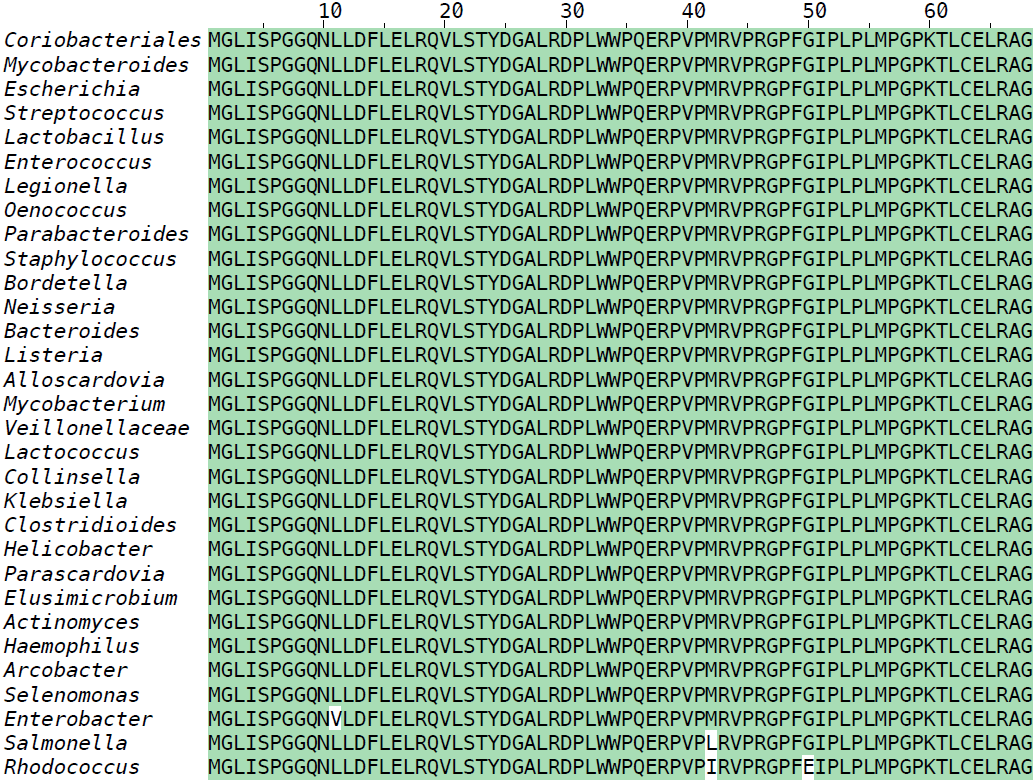
Multiple sequence alignment of 31 spurious bacterial proteins encoded on short contaminated contigs. Shown here are 31 out of 185 spurious proteins from bacterial genomes. A majority of the sequences are 100 % identical. The only differing residues are highlighted in white. This highly conserved “protein” is conserved on across different bacterial phyla, suggesting it is likely a contaminant that has been erroneously translated as part of automated annotation procedures. The respective short contigs (< 1 kb) encoding these spurious proteins align with high sequence identity and coverage to the *Ovis aries* genome.

### Contamination in the protein database NR

Conterminator can be used to analyze protein sequences. It clusters proteins [20] at 95 % sequence identity, while requiring at least 95 % sequence overlap. It reports clusters containing multiple kingdoms, using the same kingdom definition as for the nucleotide comparison. We predict that the kingdom with fewer members in the cluster is contaminated; e.g., if a cluster contains 100 proteins, and 99 represent animals while 1 represents bacteria, then the bacterial protein likely originates from a contaminated genome.

We analyzed the NCBI NR protein sequence [1] database using this procedure. We predicted 14,132 proteins to represent contaminants, out of which 7359 are also present in the Uniprot database [25]. The majority of these proteins (70.46 %) are eukaryotic, and the remaining 29.34 % are bacterial (see Supplementary Fig. 2). Over 6114 contaminant proteins originate from the phylum Arthropoda, and out of this 2401 are from the species *Trichonephila clavipes*, the golden silk orb weaver spider, which contributes overall the most contamination. We next take a closer look at this organism.

### Contaminated proteins in *Trichonephila clavipes*

The *T. clavipes* proteins identified as possible contaminants are distributed across 504 contigs with a total length of 24,152,567 residues. 163 of the contigs are longer than 20 kb. The longest contaminated contig from the assembly [26] is `MWRG01000001`, which spans 1,655,743 bases and encodes 490 proteins. We found 219 of these proteins in contaminated protein clusters, a majority of them matching the bacterium *Gemmobacter sp. YJ-T1-11*. We aligned the contig against the assembly of *Gemmobacter sp. YJ-T1-11*, and found that it is nearly 90 % covered at a 97.76 % identity (see Supplementary Fig. 3). Thus, this clearly appears to be a bacterial contig mistakenly included in the assembly of the *T. clavipes* spider.

## DISCUSSION

We present two complementary approaches to detect contamination, first using nucleotide information to detect short contaminated fragments, and second using protein-based analysis to reveal long contaminated contigs. We used a conservative approach that only considered a sequence to be a contaminant if it had a near-identical match to a species in an entirely different kingdom from the source; e.g., a bacterial sequence found in an animal genome or vice versa. Our method can efficiently detect contamination in large reference databases, and we found that a substantial fraction of the genome sequences in both GenBank and RefSeq (0.54 % and 0.34 % of entries respectively) appear to be contaminated. Contamination occurs mostly as short contigs, flanking regions on longer contigs, or regions of larger scaffolds flanked by Ns, but we also observed a few longer sequences with contamination.

Contamination can be transferred into other databases that are built from GenBank, such as the protein databases NR and Uniprot. Methods that rely on taxonomical classification, particularly metagenomics analyses, are strongly affected by cross-kingdom contamination because they often rely on subsets, e.g. microbial sequences extracted from a larger database. This makes it more difficult for such methods to detect contamination of the type reported here.

With the rapid and ongoing increase in the number of novel genomes sequenced every year, the number and variety of contaminating sequences continues to increase as well, presenting challenges for alignment-based methods to detect contamination. Conterminator’s efficiency means that it can be used routinely to detect new contamination, even on the largest databases.

## METHODS

Conterminator detects cross-kingdom alignments and predicts contamination. It builds upon existing modules of MMseqs2 [21], which it extends for use in contamination detection.

### Detection of cross-kingdom alignments

Conterminator identifies regions in genome sequences that align to genomes from other kingdoms with a minimum length of at least 100 nucleotides and a sequence identity threshold of at least 90 %. With very few exceptions, DNA sequences from different kingdoms should not be aligned at all, and sequences that match at this level of identity are strong candidates for contaminants. (Exceptions to this rule include recent horizontal gene transfer events, but these are very rare.)

Because modern sequence databases are very large, we cannot use a naive all-against-all alignment, which would entail a quadratic number of comparisons. Therefore, we used a similar strategy to Linclust [20] to reduce the computational cost to a linear number of comparisons. We reworked the algorithm to support nucleotide sequences, since Linclust was originally built to cluster protein sequences.

Conterminator first cuts all sequences into fragments of length 1000 and records their start positions. For each fragment, we extract *m* canonical *k*-mers (default *m* = 100) of length 24 with the lowest hash value and write them into an array. (We use the hash function defined in [20]; see Supplementary Figure 5.) We store the *k*-mer in 8 bytes, with the most significant bit indicating whether the *k*-mer is reversed, sequence identifiers (4 bytes), its length (2 bytes), and its position *j* in the genomic sequence (2 bytes). We sort the array by *k*-mer, length and sequence identifiers. For each *k*-mer group we assign all sequences to the longest sequence with the lowest sequence identifier *c* by overwriting their *k*-mer with the identifier of *c* and their position with the diagonal *i* − *j* respecting the strand directionality. We sort the array again with the previous criteria so that all sequences with same assignment are in a consecutive block. We write each block’s central sequence identifier, assigned sequence, strand and diagonal to hard disk while only keeping the diagonal with the most *k*-mer matches per sequence.

We perform a one-dimensional dynamic programming ungapped alignment (using the MMseqs2 command ‘`rescorediagonal --rescore-mode 2`’) on each diagonal. We assign matches a score of 2 and mismatches a score of −3 bits, and compute an E-value using ALP [27]. We compute the sequence identity by dividing the number of identical positions by the number of aligned positions. We filter out all hits that are shorter than 100 bases, or that have a sequence identity below 90 %, or that have an E-value above 10^−3^. We compute the alignment start positions by adding the start position of the fragment to the alignment coordinates (MMseqs2 command ‘offsetalignment’).

Based on the alignment, we extract the sequence intervals from *c* that are overlapped by different kingdoms. We define five “kingdoms” based on the NCBI Taxonomy [28]: (1) Bacteria & Archaea (taxonomy IDs 2 and 2157), (2) Fungi (4751), (3) Metazoa (33208), (4) Viridiplantae (33090) and (5) all other eukaryotes. We ignored sequences from Viruses (10239), unclassified sequences (12908), other sequences (28384), artificial sequences (81077), environmental samples from bacteria, archaea and eukaryotes (61964, 48479, 48510). Note that we merged Bacteria and Archaea for simplicity. We excluded viruses because some of them can integrate their genomes into other organisms, making it hard to distinguish contamination from genuine artifacts of viral integration.

### Gather all alignments by exhaustive alignment

Our detection method might miss alignments because it does not extract all *k*-mers, and because it uses 24-mers rather than shorter *k*-mers. After the previous steps, we perform an exhaustive alignment of the sequence fragments against the extracted potential contamination sequences (and their respective reverse complements) using MMseqs2. The search is performed by using the two modules `prefilter` and `rescorediagonal`. The `prefilter` program masks out low complexity regions and short tandem repeats in the potential contaminants using `tantan` [29] and detects all consecutive double 15-mers diagonal hits. We rescore the detected diagonals again with the `rescorediagonal` module, enforcing a minimal alignment length of 100 and a minimal sequence identity of 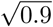. The square root of the sequence identity ensures that no pair of sequences is greater than 90 % different from each other.

### Predict contig length by find scaffolding boundaries

Genome assembly programs create scaffolds by ordering and orienting contigs using a variety of types of linking information, such as paired-end reads. A scaffold thus consists of a sequence of contigs, usually separated (in many GenBank entries) by `N`s to indicate the scaffolding boundaries. Some of the contaminants that we identified appear as short contigs in the midst of a longer scaffold, and we can identify these by finding the flanking `N`s. It is important for our contamination detection to know the real length of each contig in a scaffold. A nave approach to determining contig length would be to search for the closest `N` upstream and downstream from each alignment start and end. However, this is inefficient because many sequences contain million of bases without any `N`s. We therefore indexed all `N`’s for each sequence. We store the position of the first `N` per block in an array associated with the sequence. The `N` positions are sorted in ascending order, which enables us to perform a binary search to detect the closest `N` efficiently.

### Predict the source of contamination

A large majority of contamination occurs as small contigs (see Figure 2 in Breitweiser et al. [10]). Conterminator uses this property to help it identify contamination based on the length distribution of sequences from each kingdom. By default, it only calls a sequence a contaminant if the sequence is shorter than 20 kb, and if it aligns to a sequence in another kingdom that is longer than 20 kb.

### Predicting contamination in protein databases

Conterminator can also detect protein sequence contamination using cross-kingdom analysis. It clusters proteins using Linclust [20] with a bidirectional length overlap of 95 % and a sequence identity of 95 % (`--min-seq-id` `-c 0.95 --cov-mode 0 -a`). It reports every cluster with cross-kingdom members. For each contaminated cluster, it counts how often each kingdom occurs and reports the least abundant kingdom as the one that is contaminated. It also reports kingdoms with equal abundance, however in those cases it cannot predict the contaminated entry. Using only abundance without concern for length may lead to incorrect directionality calls.

### Data visualization

We created the Sankey plots using the `krakenuniq-report` tool from KrakenUniq [30] to create a Kraken-style report from our predicted contaminations. The visualization was done using Pavian [31] extracted as SVG and colored by Inkscape. The multiple alignment was created by MMseqs2 `result2msa` and visualized using Jalview [32]. **Software and database versions**

**TABLE 1.** List of software used in this paper. The versions for MMseqs2, KrakenUniq and Pavian are the first 6 characters of the git commit. For databases we list the date at which the data was downloaded.

| Resource | Version |
| --- | --- |
| Conterminator | fa1c80 |
| MMseqs2 [21] | 333546 |
| KrakenUniq [30] | 5c0019 |
| Pavian [31] | 81d784 |
| RefSeq [17] | July 2019 |
| GenBank [1] | Dec 2018 |
| Jalview [32] | 2.11.0 |

## Data availability

The list of contamination for Gen-Bank (`genbank.gz`) and RefSeq (`refseq.gz`) are available at: ftp://ftp.ccb.jhu.edu/pub/data/conterminator/.

## AUTHOR’S CONTRIBUTIONS

MS & SS designed research, MS developed code and performed analyses, MS & SS wrote the manuscript.

## ACKNOWLEDGEMENTS

We thank Milot Mirdita, Florian Breitwieser and Jennifer Lu for fruitful discussions. We thank the NCBI for their answering our questions timely and Aleksey Zimin for cleaning up the Turkey genome. This work was supported in part by NIH grants R35-GM130151 and R01-HG006677, and by NSF grant IOS-1744309 to SLS.

