## Supplemental Material for "Terminating contamination: large-scale search identifies more than 2,000,000 contaminated entries in GenBank"

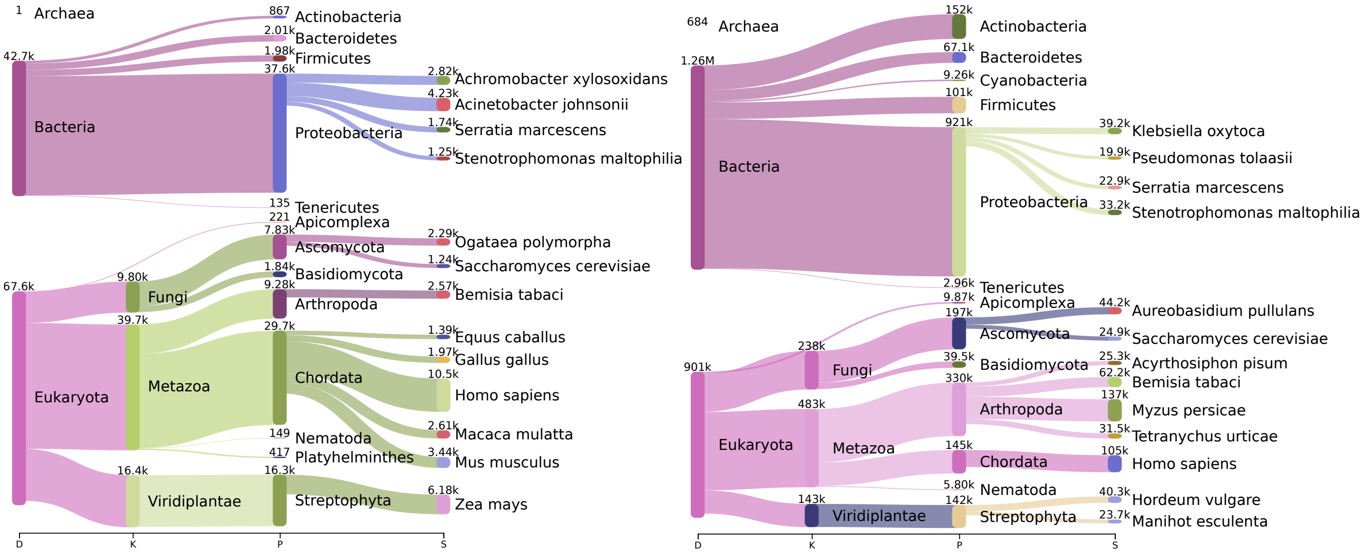


**Supplementary Figure** 1: **Sanksey plot of most contaminating species in RefSeq and GenBank. left** Sanksey plot five kingdoms: Bacteria&Archaea, Fungi, Metazoa, Viridiplantae and other Eukaryotes. **right** Distribution of contaminating species in GenBank.

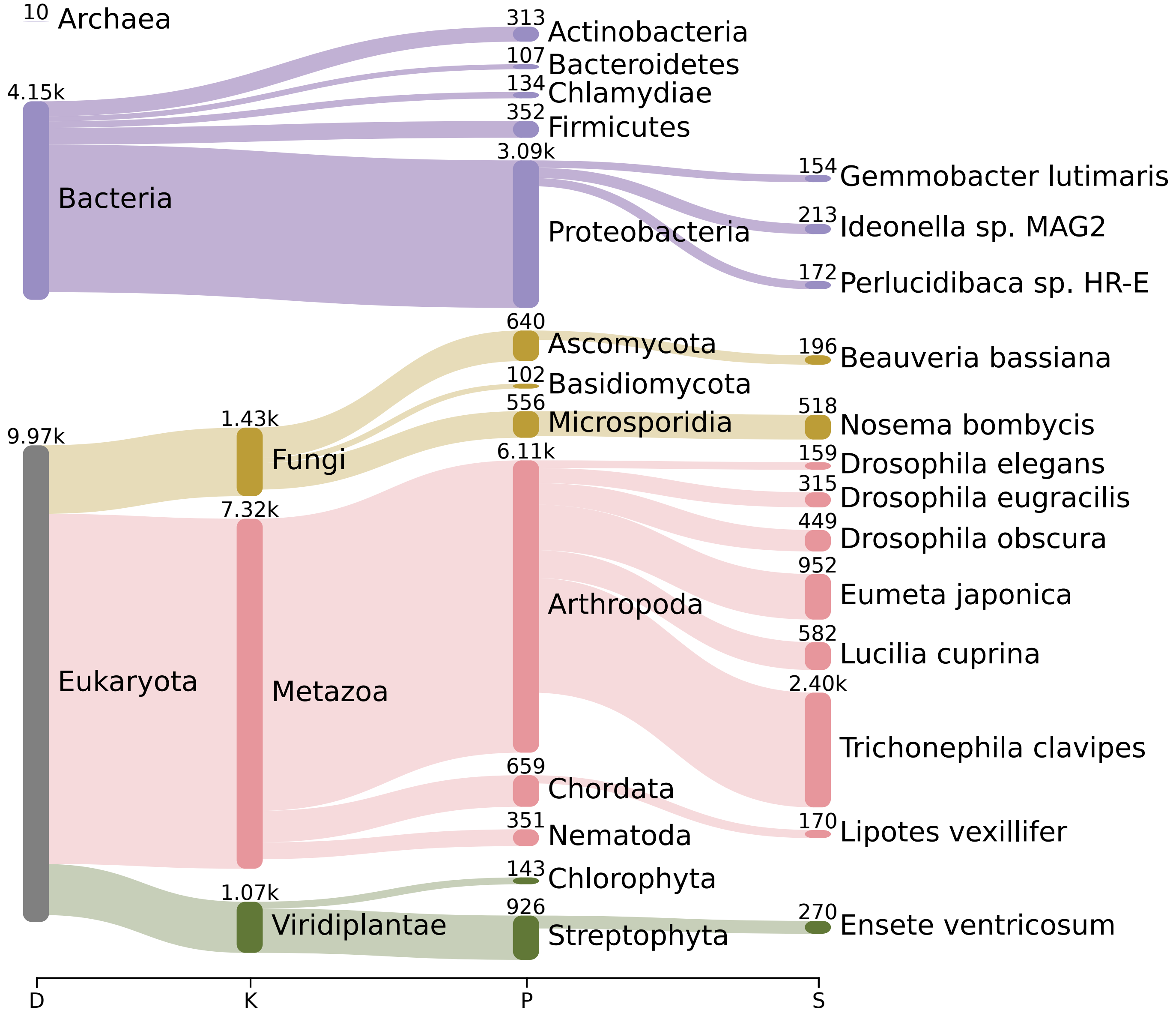


**Supplementary Figure 2: Contamination in the NR database.** Predicted contamination in NR protein database across five kingdoms.

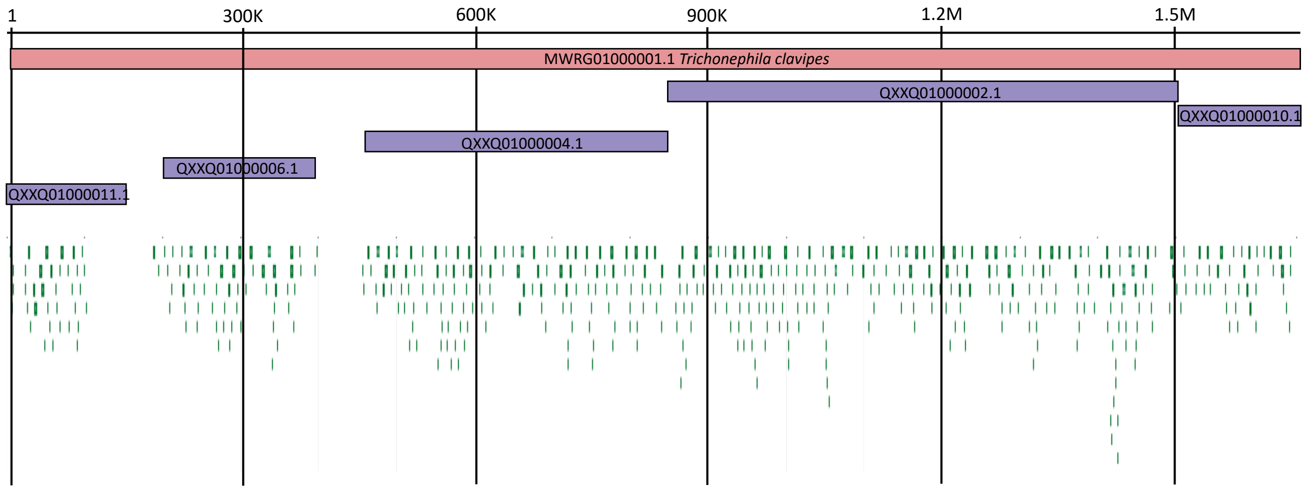


**Supplementary Figure** 3: **Longest contaminated contig of *Trichonephila clavipes*.** Alignment of the longest contaminated contig `MWRG01000001.1` of *T. clavipes* (red) and the genome *Gemmobacter sp. YJ-T1-11* (voilet). The green boxes are the annotations of *T. clavipes*, the contig encodes 490 proteins. The annotation are from the NCBI genome browser.
